# 3’-Sulfated Lewis A is a Biomarker for Metaplastic and Oncogenic Transformation of Several Gastrointestinal Epithelia

**DOI:** 10.1101/2021.03.17.435789

**Authors:** Jeffrey W. Brown, Koushik K. Das, Vasilios Kalas, Kiron M. Das, Jason C. Mills

## Abstract

**Introduction:** Multiple previous studies have shown the monoclonal antibody Das-1 (formerly called 7E_12_H_12_) specifically recognizes metaplastic and carcinomatous lesions in multiple organs of the gastrointestinal system (e.g. Barrett’s esophagus, intestinal-type metaplasia of the stomach, gastric adenocarcinoma, high-grade pancreatic intraepithelial neoplasm, and pancreatic ductal adenocarcinoma) as well as in other organs (bladder and lung carcinomas). Beyond being a useful biomarker in tissue, mAb Das-1 has recently proven to be more accurate than current paradigms for identifying cysts harboring advanced neoplasia. Though this antibody has been used extensively for clinical, basic science, and translational applications for decades, its epitope has remained elusive.

**Methods:** In this study, we chemically deglycosylated a standard source of antigen, which resulted in near complete loss of the signal as measured by western blot analysis. The epitope recognized by mAb Das-1 was determined by affinity to a comprehensive glycan array and validated by inhibition of a direct ELISA.

**Results:** The epitope recognized by mAb Das-1 is 3’-Sulfo-Lewis A (3’-Sulfo-Le^A^). 3’-Sulfo-Le^A^ is broadly reexpressed across numerous GI epithelia and elsewhere only after metaplastic and carcinomatous transformation.

**Discussion:** 3’-Sulfo-Le^A^ is a clinically important antigen that can be detected both intracellularly in tissue using immunohistochemistry and extracellularly in cyst fluid and serum by ELISA. The results open new avenues for tumorigenic risk stratification of various gastrointestinal lesions.

## INTRODUCTION

The monoclonal antibody Das-1 has been used extensively to study metaplasia and cancer in numerous tissues over the last 30 years (Fig. 1, Table 1).(1-17) Das-1 shows broad reactivity in human fetal tissue;(18) however, in adults at homeostasis, expression is primarily restricted to biliary and colonic epithelium as well as skin.(19) Despite, the absence of reactivity in normal healthy tissues of the GI foregut, the epitope is reexpressed when these tissues undergo metaplasia that increases risk for cancer and when carcinomatous transformation occurs.(1-17) Thus, the epitope recognized by Das-1 fulfills the criteria for being a true oncofetal antigen. In addition to expression in human tissues, we have recently validated the utility of Das-1 in identifying high risk pancreatic cystic lesions in a large multicenter trial, where we demonstrated that a simple ELISA for Das-1 in cyst fluid outperforms all clinical guidelines in identifying pancreatic cysts harboring malignancy.(5, 6)

**Figure 1.**
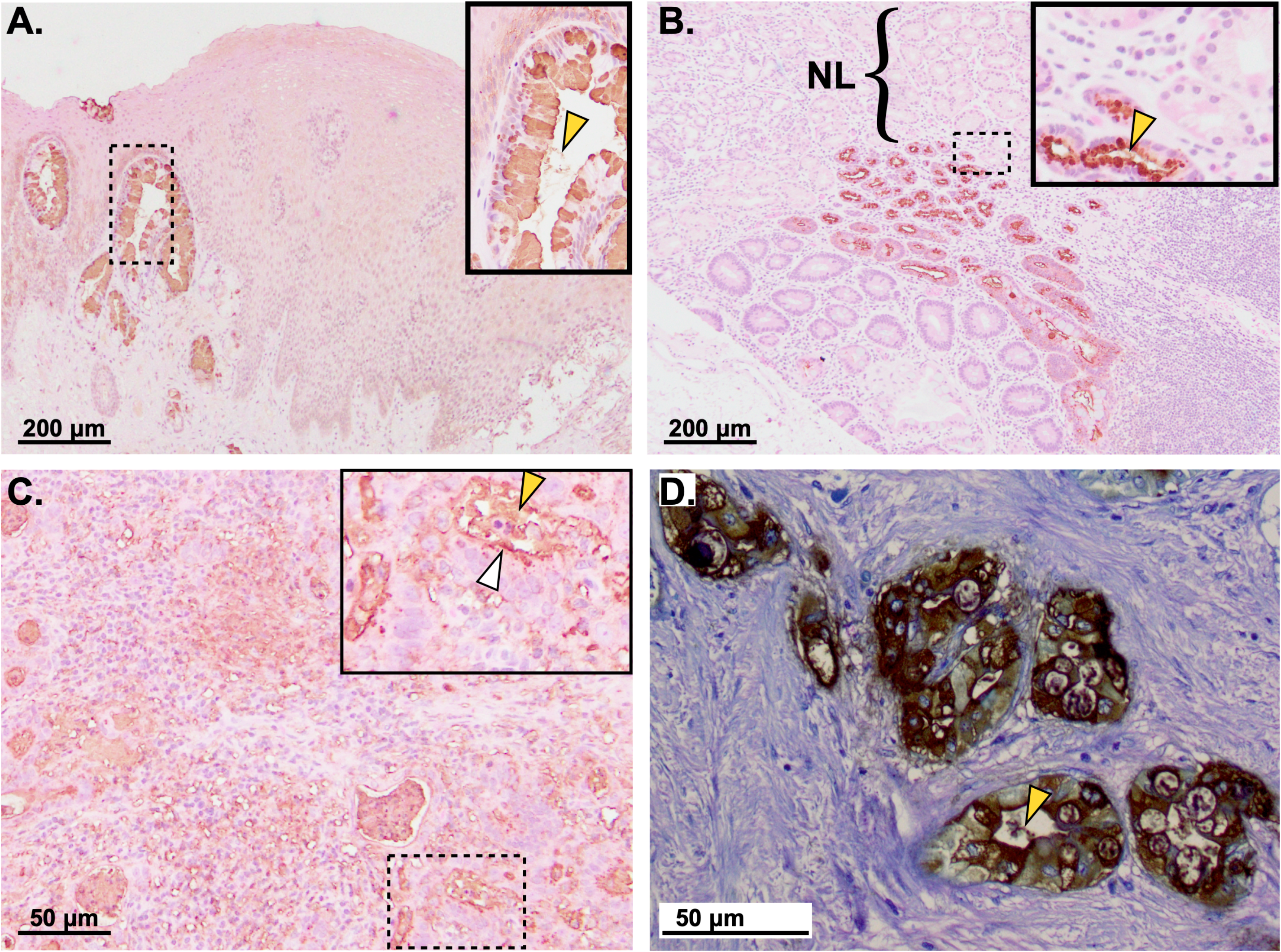
Unlike normal tissue, metaplastically and oncogenically transformed foregut tissues become reactive to mAb Das-1. Immunohistochemistry of **A**. Barrett’s Esophagus, **B**. Intestinal Metaplasia of the Stomach, **C**. Gastric Cancer, **D**. Pancreatic Ductal Adenocarcinoma. Scale bars presented in bottom left corner of each panel. Bracket labeled “NL” to highlight the absence of staining in normal stomach; contrast with incomplete intestinal-type metaplasia, which expresses 3’-Sulfo-Le^A^. Insets show higher-magnification of boxed areas. White arrowhead: Das-1 staining at a cell apex; yellow arrowhead: 3’-Sulfo-Le^A^ that has been secreted into the extracellular space.

**Table 1.** Summary of the prior literature describing Das-1 reactivity after metaplasic and/or oncogenic transformation of adult tissues that do not natively express this antigen at homeostasis.

| Tissue | Transformation | Reference |
| --- | --- | --- |
| Bladder | Cancer | Pantuck *et al.*, *J Urol* 1997; **158**: 1722-7. PMID: 9334587<br>Pantuck *et al.*, *Br J Urol* 1998; **82**: 426-30. PMID: 9772883 |
| Esophagus | Barrett's | Das *et al.*, *Ann Intern Med* 1994; **120(9)**: 753-6. PMID: 7511878<br>DeMeester *et al.*, *Am J Gastroenterol* 2002; **97(10)**: 2514-23. PMID: 12385432<br>Hahn *et al.*, *Am J Surg Pathol* 2009; **33(7)**: 1006-15. PMID: 19363439 |
| Lung | Cancer | Deshpande *et al.*, *Pathobiology* 2002; **70(6)**: 343-7. PMID: 12865630 |
| Pancreas | Cancer | Das *et al.*, *Gut* 2014; **63(10)**: 1626-34. PMID: 24277729<br>Das *et al.*, *Gastroenterology* 2019; **157(3)**: 720-30. PMID: 31175863<br>Das *et al.*, *Hum Pathol*. Accepted, in press. PMID: 33524436<br>Heidarian *et al.*, *J Am Soc Cytopathol*. Accepted, in press. PMID: 33541830 |
| Small Bowel | Adenoma | Onuma *et al.*, *Am J Gastroenterol* 2001; **96(8)**: 2480-5. PMID: 11513194 |
| Stomach | Intestinal-Type Metaplasia | Glickman *et al.*, *Am J Surg Pathol* 2001; **25(1)**: 87-94. PMID: 11145256<br>DeMeester *et al.*, *Am J Gastroenterol* 2002; **97(10)**: 2514-23. PMID: 12385432<br>Mirza *et al.*, *Gut* 2003; **52(6)**: 807-12. PMID: 12740335<br>Piazuelo *et al.*, *Mod Pathol* 2004; **17(1)**: 62-74. PMID: 14631367<br>Watari *et al.*, *Int J Cancer* 2012; **130(10)**: 2349-58. PMID: 21732341 |
| Stomach | Cancer | Mirza *et al.*, *Gut* 2003; **52(6)**: 807-12. PMID: 12740335<br>O'Connell *et al.*, *Arch Pathol Lab Med* 2005; **129(3)**: 338-47. PMID: 15737028<br>Feng *et al.*, *Exp Ther Med* 2013; **5(6)**: 1555-8. PMID: 23837030<br>Kawanaka *et al.*, *Br J Cancer* 2016; **114(1)**: 21-9. PMID: 26671747<br>Watari *et al.*, *Int J Cancer* 2012; **130(10)**: 2349-58. PMID: 21732341 |

In this study, we aim to identify the oncofetal antigen recognized by mAb Das-1 that has been used as a biomarker for high-risk metaplasia and cancer across numerous tissues in both histology as well as body fluids (serum and pancreatic cyst fluid). Here, using chemical deglycosylation, a comprehensive glycan array, and validation by inhibition of a direct ELISA, we demonstrate that the clinically important epitope of Das-1 is 3’-Sulfo-Le^A^.

## METHODS

For western blots, lyophilized antigen derived from media conditioned by the LS180 colon cancer cell line(5) was deglycosylated using anhydrous trifluoromethanesulfonic acid (TFMS) per manufacturer (Sigma-Aldrich, USA) protocol. Following the deglycosylation reaction, both control and TFMS treated antigen were diluted to the same final volume with Laemelli buffer prior to being applied to the gel. The control reaction was treated in an identical fashion (lyophilization, anisole, and pyrimidine) to the experimental condition; the only difference was TFMS was omitted.

Both the original Das-1 IgM as well as Das-1 IgG (developed from a hybridoma that had undergone spontaneous isotype switch and thus with identical *in vivo* reactivity) were assayed against a comprehensive array of 584 glycans provided by the National Center for Functional Glycomics. Briefly, the array was generated from a library of natural and synthetic mammalian glycans with amino linkers printed onto N-hydroxysuccinimide (NHS)-activated glass microscope slides forming covalent amide linkages.(20) The glycan spotting concentration was 100 μM. 6 technical replicates were performed for each antibody. Das-1 IgM was tested at 5 μg/ml and Das-1 IgG at 5 μg/ml and 50 μg/ml. Secondary anti-mouse IgM (488) and anti-Mouse IgG (488) were used at 5 μg/ml. The Das-1 IgM and IgG antibodies were provided to CFG and as a fee-for-service and the analysis against the glycan array was completed blinded.

Epitope specificity was confirmed by ELISA of wells coated with antigen (0.5 μg/well), incubated in PBS overnight at 4°C, blocked with 1% BSA, and then incubated with 2.5 μg of either Das-1 IgG or Das-1 IgM ±200 μM of the respective carbohydrate. The reactions were developed after incubation with alkaline phosphate-conjugated anti-mouse IgG or IgM and absorption at 405 nm measured. Separate assays measured effects of competitive inhibition with 0 to 200 μM 3’-Sulfo-Le^A^.

## RESULTS

Chemical deglycosylation with TFMS of a standard source of concentrated antigen recognized by Das-1 resulted in near complete loss of Das-1 binding in western blot analysis (93% and 85% as measured by IgM and IgG, respectively; Fig. 2) indicating the Das-1 epitope depended on glycans. Thus, we determined glycan specificity of the Das-1 IgM and Das-1 IgG antibodies against a comprehensive array of 584 glycans. Both Das-1 IgM and Das-1 IgG preferentially recognized Le^A^ that had been sulfated at the 3’ site of galactose (Fig. 3). The fucose is likely not essential to recognition as both antibodies recognized 3’-Sulfo-Galμ(1-3)GlcNAc as well as some disulfated glycans, albeit with lower apparent affinity. The antibodies display little-to-no affinity for the non-sulfated, sialylated, or 6’-mono-sulfated counterparts, which are listed as pertinent negatives below the highest ranked hits (Fig. 3). Recognition of the epitope was also independent of net charge, as Mannose-6-Phosphate, another negatively charged sugar, was not recognized by either antibody (Fig. 3). Relative to the IgG, the IgM isotype had similar epitope specificity but exhibited a broader range of measurable affinities (Fig. 3), as might be expected due to the greater avidity of its pentameric quaternary structure.

**Figure 2.**
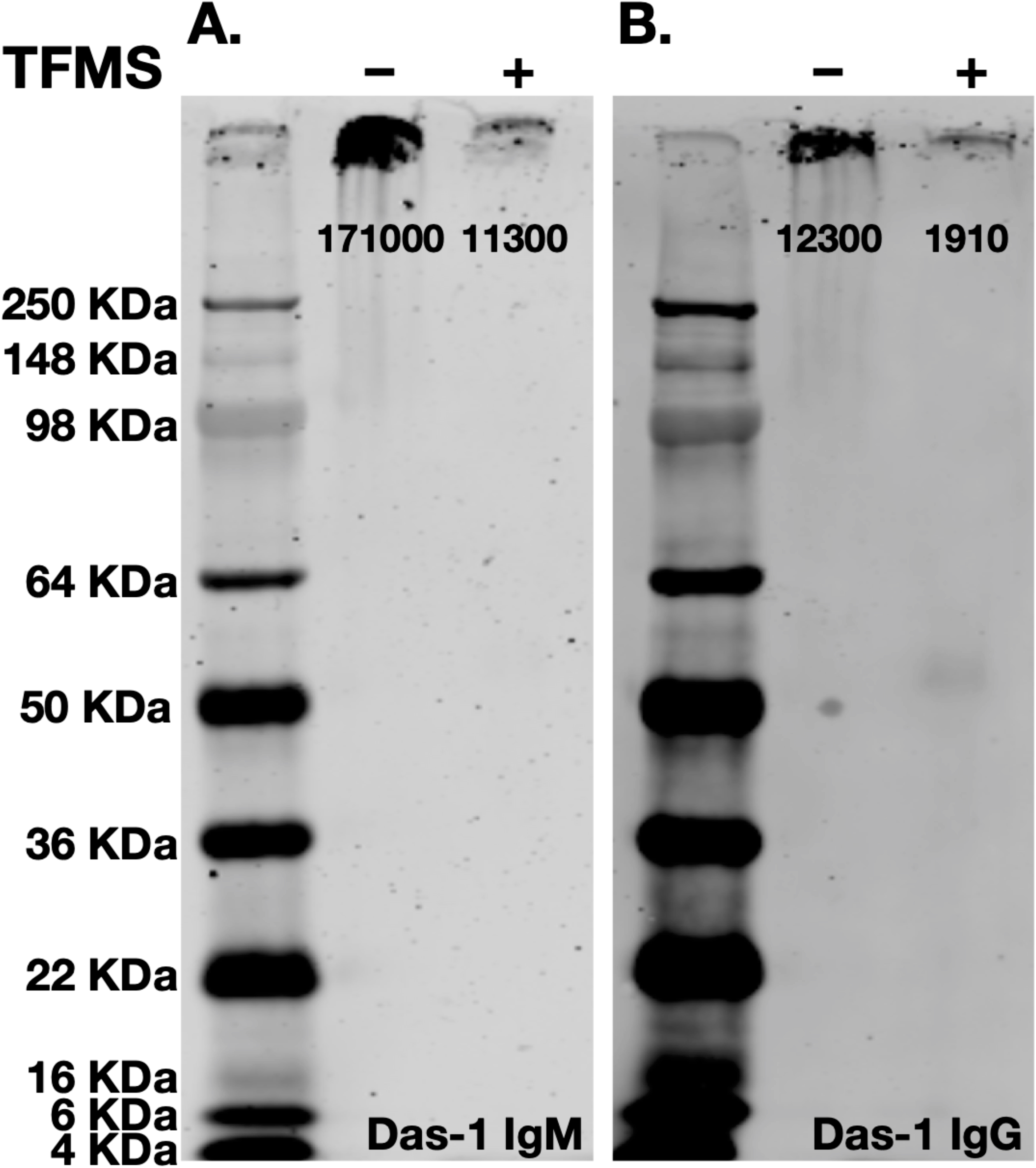
Das-1 IgG and IgM recognize a glycosylation epitope. Chemical deglycosylation of the antigen results in near complete loss of signal as measured by western blot analysis using (**A**) Das-1 IgM and (**B**) Das-1 IgG. Quantification of band intensity is presented below each band.

**Figure 3.**
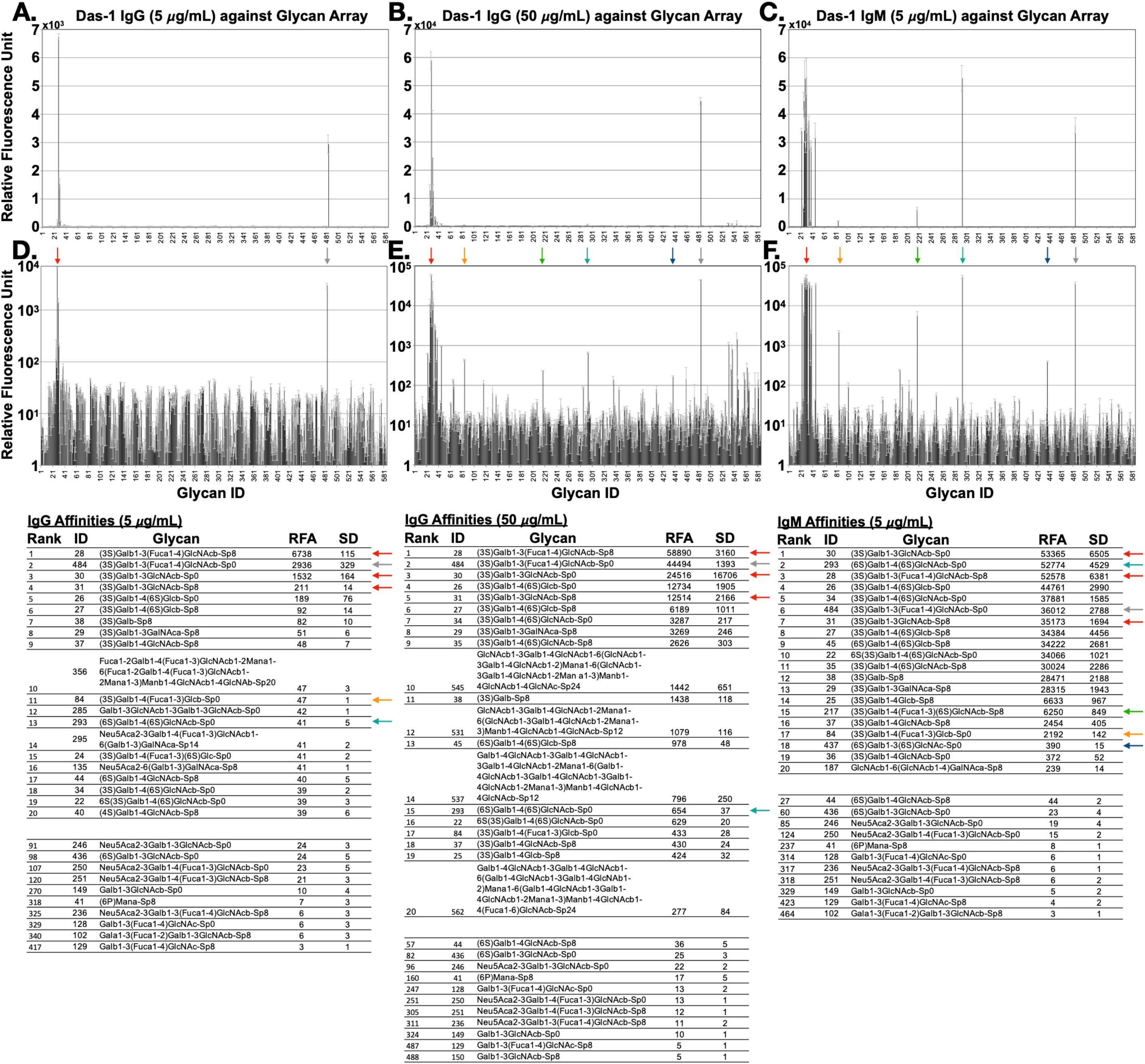
The results of the glycan arrays. Das-1 IgG (at 5 & 50 μg/mL) and Das-1 IgM (at 5 μg/mL) are plotted both linearly (A, B, C) and logarithmically (D, E, F) as the average relative fluorescence units (of 6 technical replicates) plus/minus standard deviation. The top 20 glycans for each arrays are listed and compared to non-sulfated, sialylated, 6’-monosulfated derivatives, as well as other relevant glycans. Colored arrows emphasize that Das-1 IgG and Das-1 IgM recognize the same set of glycans. The complete data sets are provided in supplemental figure 1 (5 μg/mL IgG), supplemental figure 2 (50 μg/mL IgG), and supplemental figure 3 (5 μg/mL IgM) and are available for download on the Consortium for Functional Glycomics website (www.functionalglycomics.org).

To confirm the epitope specificity, we performed a direct ELISA using Das-1 against the antigen and found that both Das-1 IgG and IgM were inhibited by 3’-Sulfo-Le^A^, in a dose-dependent manner (Fig. 4), and neither the sialylated (3’-Sialyl-Le^A^; i.e. Ca19-9) nor unsulfated adducts were able to inhibit the reaction. Despite only differing by the Galactose-GlcNAc-fucose arrangement (Fig. 4 Key), Le^X^ (type II) adducts were not able to competitively inhibit Das-1 binding in the ELISA. Further, the assay was not inhibited by sulfated galactose in the absence of the adjacent GlcNAc and fucose present in Le^A^ (Fig. 4). Thus, both the affinity and inhibitory studies presented here are consistent with 3’-Sulfo-Le^A^ being the epitope recognized by both Das-1 IgG and IgM.

**Figure 4.**
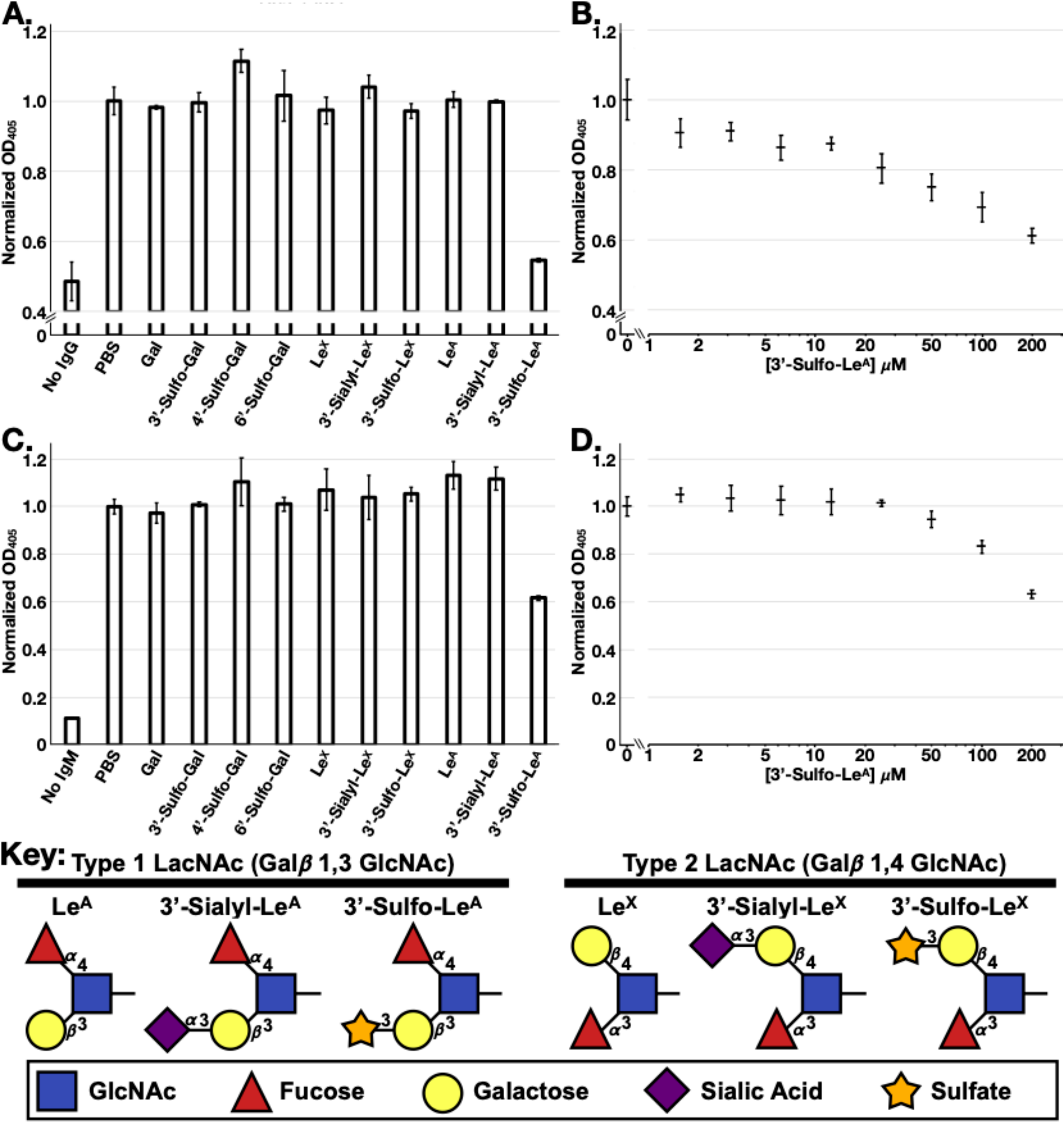
Das-1 IgG and IgM specifically recognize 3’-Sulfo-Le^A^. Direct ELISA using Das-1 IgG (**A**) or Das-1 IgM (**B**) in the absence or presence of several free glycans in solution at 200 μM. Direct ELISA using Das-1 IgG (**C**) or Das-1 IgM (**D**) against a titration series of 3’-Sulfo-Le^A^. Data reported as average ± standard deviation of three technical replicates normalized to the reaction without glycans (PBS). **Key:** Schematic diagram of the relevant Lewis antigens is provided for reference.

## DISCUSSION

Aberrant glycosylation patterns (especially acidic modification including sialylation and sulfation) have been identified in numerous types of cancer, and probing for neo-glycosylation epitopes is a common clinical practice used to (1) detect cancer, (2) monitor therapeutic response, and/or (3) evaluate for recurrence. However, typically the utility of these glycosylation epitopes is restricted to a small set of cancers (e.g. CA19-9 for pancreatic cancer). In contrast, 3’-Sulfo-Le^A^ appears to be aberrantly expressed among numerous pre-neoplastic lesions and cancers (Fig. 1, Table 1).

Sulfation is a posttranslational modification of glycans that adds a negatively charged moiety. Although, these modifications have previously been implicated in numerous cellular functions including cell adhesion and metastatic potential of tumors,(21-25) the ability to specifically detect these antigens in tissue has been stymied by the absence of commercially available lectins or antibodies.(26) Here, we demonstrate that the monoclonal antibodies Das-1 IgM and IgG specifically recognize 3’-Sulfo-Le^A^.

The progression from normal tissue to metaplasia to cancer has arguably been best described in the stomach by seminal work of Pelayo Correa and others.(27-30) The pre-cancerous state of chronic atrophic gastritis is characterized by appearance of metaplastic cells deep in the gastric glands.(31) Such Spasmolytic Polypeptide Expressing Metaplasia (SPEM) or pseudopyloric metaplasia cells express Sialyl-Le^X^ antigens that promote binding of the pro-inflammatory, carcinogenic bacteria *H. pylori*.(32, 33) Expression of these sialomucins within the columnar epithelial cells is also a defining feature of Type II, incomplete, intestinal-type metaplasia in the stomach.(34) Transition from sialylation to sulfation is the sole feature distinguishing Type II from Type III gastric intestinal metaplasia with the latter being associated with increased risk for progression to cancer.(34) The presence of sulfation in type III GIM has been well correlated with an increased risk of progressing to cancer.(34) Consistent with our data, another group has historically generated an antibody (91.9H) that recognizes 3’-Sulfo-Le^A^ and is reactive with Barrett’s esophagus(35) and GIM;(36) and phenocopies high iron diamine staining for sulfation in Barrett’s and GIM.(36) This antibody recognizes this antigen in the context of a tetra- or penta-saccharide,(37) while here we exclusively used trisaccharides and thus demonstrate that Das-1 recognizes the terminal 3’-Sulfo-Le^A^ trisaccharide and does not require other adjacent sugars.

Uncovering the diagnostically important 3’-Sulfo-Le^A^ modification has important implications. For one, new technologies for specifically detecting this glycan (*e*.*g*. mass spectroscopy (38)) may lead to even greater sensitivity in diagnosis of metaplasia and cancer at earlier stages and in a wider variety of fluids and tissues. Moreover, reproducibility of the Das-1 sandwich ELISA(5, 6) for clinical laboratory applications may be improved by using pure 3’-Sulfo-Le^A^ as a standard as opposed to the current practice for Das-1: using antigen concentrated from a colon cancer cell line.

It remains to be determined (1) *why* metaplastic and cancerous tissue of the GI foregut ubiquitously express this antigen, (2) the *necessity* of this epitope for metaplastic and oncogenic transformation, (3) what proteins or lipids carry this epitope, and (4) the molecular mechanism by which this epitope is released in pancreatic cyst fluid(5, 6) as well as in the serum of individuals with cancer.(38, 39) If the cellular processes annotated by these 3’-Sulfo-Le^A^ antigens confer a proliferative or survival advantage to cancer then specifically inhibiting the sulfation reaction may provide a novel therapeutic strategy for these lethal cellular transformations.

## ACKNOWLEDGEMENTS

Jeffrey W. Brown is supported by the Department of Defense, through the PRCRP program under Award No. W81XWH-20-1-0630, NIH T32 DK007130-42, the Digestive Disease Research Core Centers Pilot and Feasibility Grant as part of P30 DK052574, and the American Gastroenterological Association AGA2021-5101. Jason C. Mills is supported by the National Institute of Diabetes and Digestive and Kidney Diseases (R21DK111369, R01DK094989, R01DK105129, R01DK110406), the Alvin J. Siteman Cancer Center-Barnes Jewish Foundation Cancer Frontier Fund, The National Institutes of Health National Cancer Institute (P30 CA09182, R01CA239645, R01CA246208), and the BETRNet (U54CA163060). The National Center for Functional Glycomics Glycan Array resource and much appreciated assistance with the analysis was supported by R24 GM098791, R24 GM137763, and P41 GM103694. V.K. was supported by the Medical Scientist Training Program Training grant T32 GM07200. Development of mAb Das-1 was supported in part by research grants NIDDK, R01 DK47673 and R01 DK63618 to KMD.

## AUTHOR CONTRIBUTIONS

JWB, KKD, VK, JCM: Conceived of the project, funded the research, performed experiments, analyzed data, and wrote the manuscript KMD: Provided essential reagents, funded the research, and edited the manuscript.

## STATEMENT OF ETHICAL ASSURANCE

JWB & JCM is the guarantor of this work and, as such, had full access to all of the data in the study and takes responsibility for the integrity of the data and the accuracy of the data analysis.

---

Supplemental Dataset 1. Complete glycan array dataset of Das-1 IgG (5 ug/uL) against the CFG glycan Array. This dataset will also be released for public download at www.functionalglycomics.org.

Supplemental Dataset 2. Complete glycan array dataset of Das-1 IgG (50 ug/uL) against the CFG glycan Array. This dataset will also be released for public download at www.functionalglycomics.org.

Supplemental Dataset 3. Complete glycan array dataset of Das-1 IgM (5 ug/uL) against the CFG glycan Array. This dataset will also be released for public download at www.functionalglycomics.org.

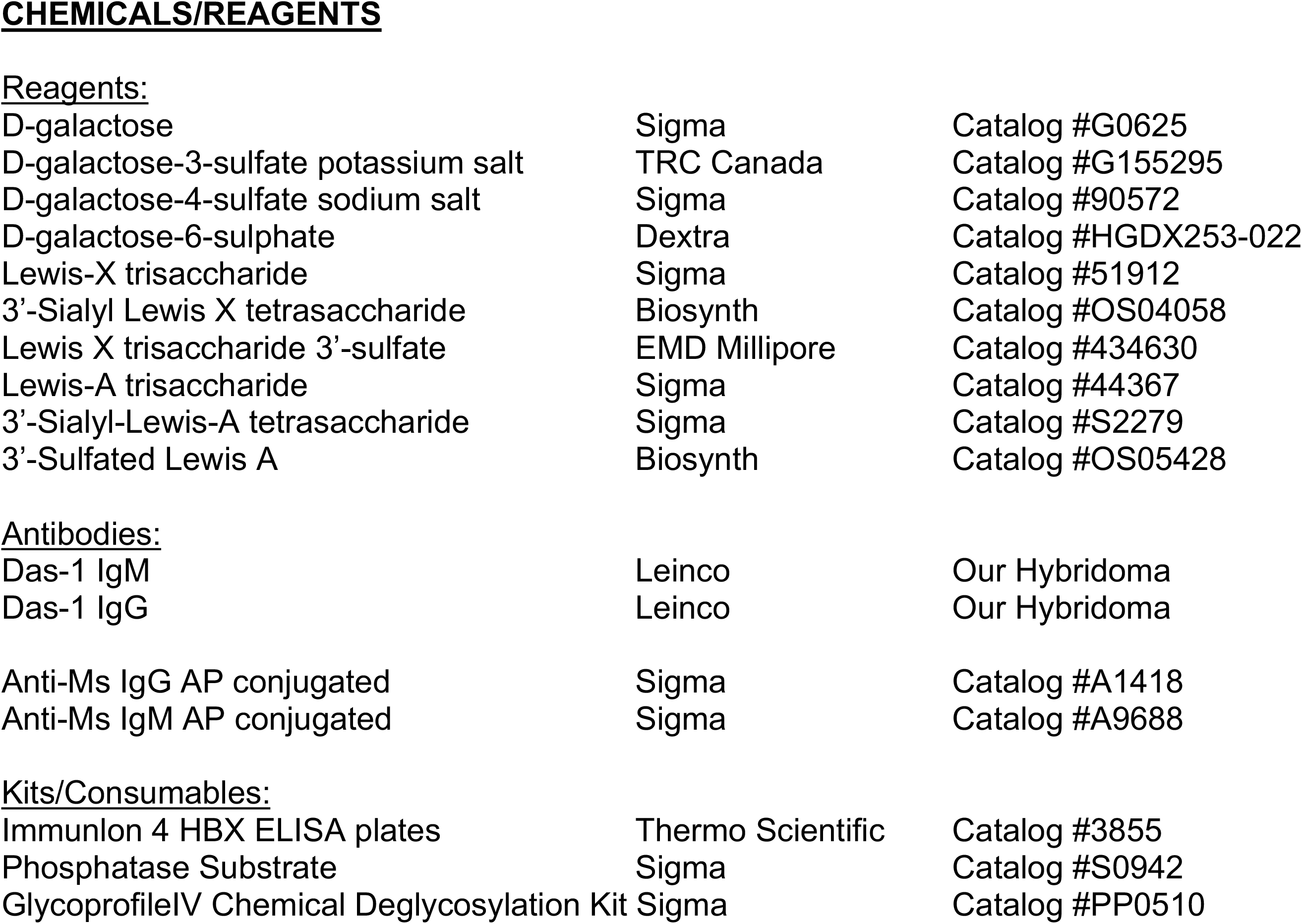

## Abbreviations used in this paper

BSA: Bovine Serum Albumin
ELISA: Enzyme Linked Immunosorbent Assay
Gal: Galactose
GlcNAc: N-Acetylglucosamine
PBS: Phosphate Buffered Saline
Gal: Galactose
Le^A^: Lewis A
Le^X^: Lewis X
TFMS: Trifluoromethanesulfonic Acid

